# Distinguishing host responses, extensive viral dissemination and long-term viral RNA persistence in domestic sheep experimentally infected with Crimean-Congo hemorrhagic fever virus Kosovo Hoti

**DOI:** 10.1101/2023.08.04.552053

**Authors:** Hongzhao Li, Mathieu Pinette, Greg Smith, Melissa Goolia, Bradley S Pickering

**Affiliations:** National Centre for Foreign Animal Disease, Canadian Food Inspection Agency, Winnipeg, Manitoba, Canada; Department of Medical Microbiology and Infectious Diseases, College of Medicine, Faculty of Health Sciences, University of Manitoba, Winnipeg, Manitoba, Canada

**Keywords:** Crimean-Congo hemorrhagic fever virus, Kosovo Hoti, sheep, viremia, host response, antibody, cytokine, virus dissemination, virus shedding, virus persistence

## Abstract

Crimean-Congo hemorrhagic fever orthonairovirus (CCHFV) is a tick-borne, biosafety level 4 pathogen that often causes a severe hemorrhagic disease in humans (CCHF) with high case fatality rates. The virus is believed to be maintained in a tick-vertebrate-tick ecological cycle involving numerous wild and domestic animal species, however the biology of CCHFV infection in these animals remains poorly understood. Here, we challenge domestic sheep with CCHFV Kosovo Hoti, a highly pathogenic clinical isolate increasingly utilized in current research. In the absence of prominent clinical signs, the infection leads to an acute viremia and coinciding viral shedding, high fever and markers for potential impairment in liver and kidney functions. A number of host responses distinguish the subclinical infection in sheep versus fatal infection in humans. These include an early reduction of neutrophil recruitment and its chemoattractant, IL-8, in the blood stream of infected sheep, whereas neutrophil infiltration and elevated IL-8 are features of fatal CCHFV infections reported in immunodeficient mice and humans. Several inflammatory cytokines that correlate with poor disease outcomes in humans and have potential to cause vascular dysfunction, a primary hallmark of severe CCHF, are down-regulated or restricted from increasing in sheep. Of particular interest, the detection of CCHFV RNA in a variety of sheep tissues long after the acute phase of infection indicates a widespread viral dissemination in the host and suggests a potentially long-term persisting impact of CCHFV infection. Consistent with this, antibody responses exhibit features reminiscent of recurring antigenic boost, and a prolonged fever or late fever spike correlates with high levels of viral RNA persistence. These findings reveal previously unrecognized aspects of CCHFV biology in animals and highlight the need for extended experimental infection studies.

**Author summary:** Crimean-Congo hemorrhagic fever orthonairovirus (CCHFV) is a tick-borne virus with potential to cause a fatal hemorrhagic disease in humans. Many wild and domestic animals such as sheep are believed to serve as intermediate hosts that amplify and transmit the virus without developing overt disease. However, the biology of CCHFV infection in animals remains to be better understood through new experimental infection research. Here, we characterize the infection of sheep with a highly pathogenic (to humans) CCHFV clinical isolate. This work confirms early studies indicating that CCHFV infection in animals does not lead to prominent signs of disease despite a short period of viral accumulation in the blood. Importantly, we identify host responses that distinguish the lack of disease in sheep versus the fatal disease in humans. Sheep are able to restrict several immune factors that potentially play a damaging role toward poor disease outcomes. Furthermore, we provide pioneering findings of widespread CCHFV dissemination and persistent presence of CCHFV genetic material in tissues of animal hosts that do not develop major disease. These new data are anticipated to inform medical countermeasure development and guide public health measures, with considerations of potential long-term impact of CCHFV on human and animal health.

## Introduction

CCHFV is a biosafety level 4 pathogen characterized by high case fatality rates (CFR) in humans ranging from 5% to 30% but up to 80% in some outbreaks [1–4] and an absence of internationally licensed vaccines or therapeutics. The virus has been listed by the World Health Organization as a top priority pathogen of epidemic and pandemic potential in urgent need for research and development [5, 6]. Belonging to the genus *Orthonairovirus*, family *Nairoviridae*, order *Bunyvirales* [7], CCHFV is an enveloped virus with an RNA genome consisting of the small (S), medium (M) and large (L) segments, which encode the nucleoprotein, glycoprotein precursor and RNA-dependent RNA polymerase, respectively [1]. Ixodid (hard-body) ticks from the genus *Hyalomma* are the principle reservoir and vector for the virus and human infections most often occur through tick exposure [1]. CCHFV has a widespread geographic distribution, involving Asia (from Western China to the Middle East and Turkey), Europe (Eastern and Southeastern countries) and Africa (majority of the continent). This vast geographic range reflects that tick hosts tolerate broadly diverse environments [1, 8]. There have been emerging signs that CCHFV is expanding into new territories such as Western Europe, which may be facilitated by emergence of favorable ecological environment driven by climate change and introduction of infected ticks by migratory birds or livestock trade [1, 4, 8–13]. In recent years there has been a trend of rising global incidence of CCHFV infections, with greatly increasing case numbers reported from several major endemic countries, demonstrating an imminent public health impact of this re-emerging virus [14–22].

In humans CCHFV infection can result in a range of disease outcomes [1, 23–25]. Most cases are asymptomatic or mild with non-specific symptoms, such as fever, headache, myalgia, dizziness, back and abdominal pains, nausea, vomiting and diarrhea. However, some cases quickly progress to a severe, often fatal, hemorrhagic fever disease (CCHF), characterized typically by vascular dysfunction, hemorrhagic manifestations, multi-organ failure (including cerebral, liver, and kidney failure and cardiac and pulmonary insufficiency), shock and death [1, 24–26]. Some CCHF survivors were reported to experience sequelae that may persist for longer than a year [23]. A number of laboratory findings are common correlates or predictors of poor outcomes, including high viral load (viremia), elevated serum levels of liver-associated enzymes, aspartate and alanine aminotransferases (AST and ALT), disseminated intravascular coagulation (DIC), thrombocytopenia, prolonged clotting times, absent or weak antibody response and increased serum levels of inflammatory cytokines and chemokines (hereafter both referred to as “cytokines” for simplicity) [1, 23–25, 27, 28].

In wild and domestic vertebrate species, natural CCHFV infection does not appear to produce prominent disease. Humans are the only known susceptible host to experience severe CCHF disease. A few exceptions exist in laboratory animals. Newborn mice, immunodeficient mice or hamsters, humanized mice and most recently cynomolgus macaques (with variable results) have demonstrated CCHF-like disease [1, 29–33]. For simplicity, other than these CCHF-susceptible laboratory animals, we refer to wild and domestic vertebrate species as “animals” hereafter. Field data on CCHFV infection in animals are primarily restricted to seroepidemiological surveys where CCHFV antibodies have been detected in a wide range of hosts, while the virus was isolated in a small number of cases [29, 34–39]. Like other large herbivores, domestic ruminants (sheep, cattle and goats) support and carry large numbers of infected ticks and exhibit a high prevalence of CCHFV antibodies [1, 40]. Sheep have been epidemiologically linked to human cases of CCHFV infection on a number of occasions [1, 40–45].

In experimental CCHFV infection studies, animals in general manifest no overt clinical disease, but often develop a transient viremia, and viral transmission from infected animals to ticks have been observed in a number of cases. Birds do not develop detectable viremia, with a few exceptions; however, aviremic birds were reported to transmit the virus to ticks [46]. Due to the lack of obvious disease in animals, CCHFV is not considered to have direct economic impact on livestock production. However, animals carrying viremia in addition to infected ticks represent a source of viral transmission. This has led to restrictive public health advices or measures, such as the ban of livestock transportation or slaughtering, or the closure of farms, which can cause economic impact indirectly [47–52].

The experimental infection studies, together with serosurveys, have largely established a tick-vertebrate-tick ecological cycle for CCHFV maintenance and transmission. In addition to their role in feeding, carrying and transporting ticks, a variety of animal species serve as viral amplification hosts. However, numerous gaps remain in the research of CCHFV infection in animals. Past experimental studies were reported during the period of 1945 – 1978, many in the Russian-language literature, with only six published more recently (1980s – 1990s) [1, 23, 29]. It has been noted that findings from these studies remain to be validated using modern laboratory methods [1]. Conclusions from these studies may also have been confounded by variations in inoculum source, which was based on viral strains of unclear pathogenicity or poor characterization [29]. Data from experimental infections of animals have so far been limited to the assessment of viremia and, in some studies, antibody responses, without in-depth clinical and biological findings [29]. Host responses associated with disease control, viral shedding and viral dissemination and persistence in tissues of animals are notable gaps in CCHFV knowledge [29, 53]. Novel insights gained by addressing these questions is anticipated to identify candidate targets for medical countermeasure development and provide critical guidance for public health education and interventions in One Health management.

In the current study, we conducted experimental infections of domestic sheep using the Kosovo Hoti strain of CCHFV. Originally isolated from a fatal human case from an endemic region with high CFR [54], the strain represents high human pathogenicity and is increasingly used in current research toward CCHFV medical countermeasures as well as in pathogenicity studies [32, 55–58]. Our detailed analysis of virological, clinical and immunological parameters associated with CCHFV infection in sheep identified differential host responses contrasting those known for fatal infections in humans and immunodeficient mouse models. Further, this work provided evidence of viral shedding concerning different routes, widespread viral dissemination in the host and long-term viral RNA persistence in tissues.

## Materials and Methods

### Ethics and biosafety approval statement

All animal work was approved by the Animal Care Committee of the Canadian Science Centre for Human and Animal Health, under the animal user document number C-21-001. The study was designed and performed by strictly following the Canadian Council on Animal Care guidelines and Russell and Burch’s 3Rs Principles [59, 60]. Animal housing, care, environmental enrichment and humane treatments during experimental procedures were conducted as previously described [61]. All invasive procedures, including viral inoculation, sample collection and euthanasia, were performed under isoflurane anesthesia. At the end of the study, animals were euthanized by intravenous administration of a commercial sodium pentobarbital solution. All experiments involving infectious CCHFV were conducted in the containment level 4 laboratory at the National Centre for Foreign Animal Diseases, Canadian Food Inspection Agency, following the institutional standard operating procedures.

### Virus, cells and animals

CCHFV Kosovo Hoti (GenBank accession numbers DQ133507, EU037902 and EU044832 for the S, M and L segments, respectively; passage 7) was acquired from the European Virus Archive – Global (https://www.european-virus-archive.com/virus/crimean-congo-hemorrhagic-fever-virus-strain-kosovo-hoti; Ref-SKU: 007V-02504) [32, 57]. The original viral stock was subjected to one passage on SW-13 cells (ATCC, CCL-105) [62–64] to generate a working stock (passage 8) used for animal inoculation. The viral titer was determined on SW-13 cells using a TCID50 (50% tissue culture infectious dose) method. Detailed methods of viral production and titration were described previously, based on the CO_2_^+^ culture condition [64]. Four domestic sheep (*Ovis aries*) of the Rideau-Arcott breed, male and approximately two months old, were sourced from a high health status farm in Manitoba, Canada and used for infection with CCHFV. These animals were named Sheep 21-01, 21-02, 21-03 and 21-04. 10 uninfected sheep remaining in the same farm flock as the four sheep for infection originated from, with the same/similar biological characteristics (breed, sex and age), were involved in blood and swab sampling to provide additional control samples, but were not challenged with the virus or sacrificed. These uninfected sheep were only subjected to sampling once and no complications would be expected to occur following this sampling event.

### Study design

The four animals for experimental infection were injected with CCHFV Kosovo Hoti via both the subcutaneous and intravenous routes at 1.2 × 10^6^ TCID50 per route. A daily visual check of apparent wellbeing was performed between −1 and 34 days post infection (DPI), the baseline and study endpoint days, respectively. Recording of clinical signs of illness and collection of blood and swabs (nasal, oral and rectal) was conducted daily on −1, 1-10, 14, 21, 28 and 34 DPI, except that during 1-10 DPI, two animals were subject to blood and swab sampling every other day (Sheep 21-01 and 21-03 on odd number DPI, whereas 21-02 and 21-04 on even number DPI). The alternate day sampling was initiated to reduce the distress on the animals so as not to subject them to every day sampling for 10 consecutive days. On 34 DPI, after routine sampling all animals were euthanized and the following organs/tissues were collected: skeletal muscle - right quadriceps, inguinal lymph nodes, gastrohepatic lymph nodes, liver, spleen, mesenteric lymph nodes, ileum, adrenal gland, kidney, lung - right middle lobe, heart, tracheobronchial lymph nodes, deep cervical lymph nodes, cerebrum, cerebellum, hypothalamus, testicle, lung - left cranial lobe, additional lung tissue and cerebrospinal fluid.

To address the clinical and biological responses to CCHFV infection in the four challenged sheep, changes were analyzed between the baseline (before infection) and subsequent time point of interest (after infection), or in other words, between the “baseline control group” and “infected group”. The smallest sample size followed the 3Rs principles and was calculated using the UCSF Sample Size Calculators for designing clinical research [65]. This was based on a type 1 error, α (two-tailed) = 0.05, and a type 2 error, β = 0.2, the proportion of subjects that are in group 0 (unexposed), q0=0.5, the proportion of subjects that are in group 1 (exposed), q1=0.5, risk in group 0 (baseline risk), P0 = 0.001 and risk in group 1 (exposed), P1 = 0.9. The estimated results indicated no false positives or false negatives. To enhance statistical power, an enlarged negative control group (NC) of 14 sheep was formed by combining the four sheep for infection at the baseline state with the 10 additional uninfected sheep. Thus, an extended analysis was performed comparing samples from the enlarged NC group and samples from the infected sheep group.

### Statistical analysis

Significance of difference was determined by *Student’s paired t* test between the baseline control group and infected group and by *Student’s unpaired t* test between the enlarged NC group and infected group, using the GraphPad Prism software. A *p* value less than 0.05 was defined as significant.

### Viral RNA analysis

Sample inactivation was performed by mixing rigorously 175 µl sodium citrate-treated blood with TRIzol LS Reagent (ThermoFisher Scientific, 10296028), or by mixing 70 µl swab elute or tissue homogenates with 630 µl TriPure Isolation Reagent (Sigma–Aldrich, 11667165001). Purification and RT-qPCR quantification of CCHFV RNA were conducted as previously described [64].

### Hematology, chemistry and coagulation times

Hematological analysis of blood cell counts and characteristics was performed using a VetScan HM5 hematology analyzer (Zoetis, https://www.zoetis.com/products-and-science/products/vetscan-hm5) on EDTA-treated whole blood. Blood chemistry including markers for liver and kidney diseases was analyzed on a VetScan VS2 chemistry analyzer (Zoetis, https://www.zoetis.com/products-and-science/products/vetscan-vs2) using the Preventative Care Profile Plus rotor with lithium heparin-treated whole blood. Citrated whole blood was used to test for changes in coagulation times using Coag Dx Analyzer (IDEXX, https://www.idexx.com/en/veterinary/analyzers/coag-dx-analyzer/) with Citrated Blood PT (IDEXX, 99-13884) and Citrated Blood aPTT (IDEXX, 99-13885) cartridges. Compliance with the manufacturer’s instructions was ensured for all the procedures. Details of the measured parameters and resulting data are provided in Table S1.

### Indirect IgG ELISA

Nunc MaxiSorp flat bottom 96-well ELISA plates (Sigma-Aldrich, M9410) were coated with 50 ng/well of recombinant CCHFV Gn protein (Creative Diagnostics, DAGF-200) or nucleoprotein (Creative Diagnostics, DAGA-3109) in 50 µl 0.06 M carbonate/bicarbonate buffer with pH 9.6 at 4 °C overnight. Coated plates were washed three times using 0.01 M PBS containing 0.05% Tween20 with pH 7.2 (PBS-T). The wash was similarly carried out between each of the following steps. The plates were blocked with 5% skim milk in PBS-T (blocking buffer) for 1 hour (hr) at 37°C. Sera diluted at 1:500 in blocking buffer was added at 100 µl/ well and incubated at 37°C for 1 hr. The secondary antibody, Donkey anti-Sheep IgG (H+L) Secondary Antibody, HRP (ThermoFisher Scientific, A16041) was diluted at 1:2000 in blocking buffer and used at 100 µl/well with incubation at 37°C for 1 hr. For enzymatic color development, the Pierce TMB Substrate Kit (ThermoFisher Scientific, 34021) was used following the supplier’s protocol. Optical density values were read on a BioTek Epoch Microplate Spectrophotometer (Agilent).

### Virus neutralization test

Neutralizing antibody titers in serum samples were determined by a plaque reduction neutralization test (PRNT) against CCHFV. Serum samples were heat inactivated at 56 °C for 30 minutes (min), serially fivefold diluted and incubated with virus at 37 °C for 1 hr. Each serum-virus mixture was then applied to an antibody staining-based plaque assay described previously [64]. The highest dilution fold with > 70% reduction in plaque counts compared to uninfected control sera was defined as the neutralization titer.

### Luminex cytokine assay

The MILLIPLEX Ovine Cytokine/Chemokine Panel 1 Premixed 14-plex - Immunology Multiplex Assay (Millipore Sigma, SCYT-91K-PX14) was used to quantify 14 cytokines in sheep serum samples following the manufacturer’s instructions. These cytokines include interferon gamma (IFN-γ), interferon gamma-induced protein 10 (IP-10, also known as CXCL10), interleukin-1 alpha (IL-1α), macrophage inflammatory protein-1 alpha (MIP-1α, CCL3), IL-8 (CXCL8), vascular endothelial growth factor A (VEGF-A), IL-1β, IL-17A, MIP-1β (CCL4), tumor necrosis factor alpha (TNF-α), IL-6, IL-10, IL-4 and interleukin 36 receptor antagonist (IL-36Ra). Data were generated based on a 1:2 sample dilution for all cytokines except 1:8 for IL-8 and IP-10. The assay plate was run on a Luminex MagPix system using the xPONENT software of version 4.2.1324 (Luminex). Conversion of median fluorescent intensity (MFI) values to pg/ml was performed using an unweighted 5-parameter logistic method in xPONENT when all sample values for a cytokine fell within the calculation range of the standard curve.

### Tissue homogenization

Tissues in Precellys bead mill tubes (Bertin, https://en.esbe.com) were mixed with DMEM (Wisent, 319-005-CL) at a 10% weight-to-total volume ratio and processed on a Minilys tissue homogenizer (Bertin, https://www.bertin-instruments.com) at maximum speed for 45 seconds. The homogenates were clarified as supernatants after centrifugation at 3000 g for 15 min (4 °C). For antibiotic treatment, homogenates were mixed with a 100× glutamine-antibiotic stock solution [66] at a 9:1 volume ratio (final antibiotic concentration at 10×) and incubated for 30 min at room temperature. The antibiotic-treated homogenates were further clarified by centrifugation as above. To make the 100× glutamine-antibiotic stock solution, a Solution A was prepared by dissolving 2.92 grams of L-glutamine (Sigma, G8540-25G) in 50 ml of sterile water and sterilized by passing through a 0.22 µm filter. A Solution B was prepared by dissolving 1,000,000 international units penicillin G sodium salt (Sigma, P3032-10MU), 1 gram streptomycin sulfate (Sigma, S9137-25G), 500,000 units nystatin (Sigma, N6261-500KU), 150,000 units polymyxin B sulfate (Sigma, P4932-1MU) and 1 gram active kanamycin monosulfate (Sigma, K1377-5G) in 5 to 10 ml of sterile water each and pooling these individual solutions together. Solution A and Solution B were then aseptically mixed and the total volume was brought to 100 ml with sterile water. The resulting 100× glutamine-antibiotic stock solution was aliquoted and stored at −20 °C.

### Virus isolation

For virus isolation from blood, 100 µl/ well of sodium citrate-treated blood, undiluted or diluted at 1:100 or 1:1000 with DMEM (Wisent, 319-005-CL), was added onto SW-13 cells (approximately 90% confluent) cultured in 48 well plates following removal of old media. Viral adsorption was allowed for 1 hr. The inoculum was then removed and wells were washed two times with 300 µl maintenance media (DMEM + 2% FBS + 100 IU/ml penicillin and 100 µg/ml streptomycin), followed by addition of 600 µl fresh maintenance media. After incubation for 7 days, the growth of infectious CCHFV was determined by both the observation of cytopathic effect (CPE) in the cell culture [64] and the detection of viral RNA genome amplification with RT-qPCR as described above. It should be noted that no blood toxicity to SW-13 cells was observed; however, a 1:10 blood dilution in DMEM led to the formation of large clots, making the proper transfer of samples impossible. For virus isolation from tissues, untreated and antibiotic-treated tissue homogenates (described above), undiluted or diluted at 1:10 in DMEM, were subjected to the same virus isolation procedure as with blood. The exception, however, was that following the observation of no CPE on 5 DPI of the tissue viral isolation, the culture supernatants were transferred onto fresh SW-13 cells for a 7-day, second round of virus isolation test.

## Results and discussion

### Susceptibility of lambs to CCHFV infection

Our study design included factors previously assessed for correlation with seroprevalence of CCHFV antibodies in both animals and humans. While sex showed no effect in sheep [67], increasing age was associated with higher antibody prevalence in sheep, other domestic animals and humans [43, 45, 67–75]. Age may reflect the amount of exposure of livestock or humans to infected ticks [71]. Alternatively, cases in younger aged animals may be linked to higher resistance to infection. In newborn mice, however, younger age appears to confer higher susceptibility to CCHF disease. Taking the potential effect of age into account, we focused this study on characterizing CCHFV infection in sheep of younger age. Four male lambs at approximately two months of age, identified as Sheep 21-01, 21-02, 21-03 and 21-04, respectively, were each challenged with CCHFV Kosovo Hoti through both the subcutaneous and intravenous routes (1.2 × 10^6^ TCID50 per route). Animals were monitored for clinical signs daily and sampled on alternate days or weekly. Viremia (or viral load), as measured by viral RNA in whole blood, was detected in all animals, at time points ranging from 2 DPI to 6 DPI, with peak levels observed around 5 DPI and followed by a quick fall to undetectable levels (Figure 1). We performed virus isolation on blood samples covering time points from −1 DPI (baseline) to 8 DPI. Infectious virus was isolated in all animals, from the time points within the viremic period (2 DPI – 6 DPI) (Figure 1). These results demonstrate that lambs are susceptible to CCHFV infection. This also confirms the similar susceptibility observed by Zarubinsky *et al.* (1976) in lambs and others in adult sheep [29, 76–79].

**Figure 1.**
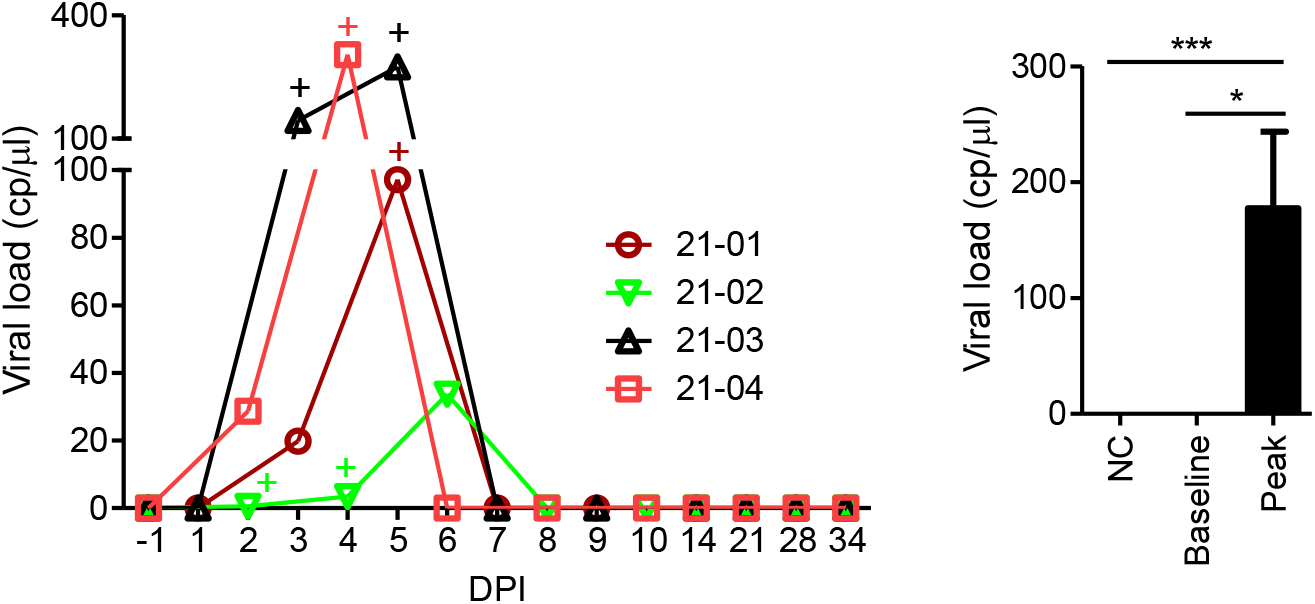
Viremia in CCHFV-infected sheep. Viremia, indicated as viral load, was measured by viral RNA in whole blood. Line graph shows the viral load time course for each experimentally infected animal. The “+” symbol marks the time points where infectious virus was isolated. Bar graph compares viral loads at their peak in the infected animals against those at the baseline (−1 DPI) or those in 14 uninfected, negative control (NC) animals. The induction of viremia by CCHFV infection was found statistically significant: \**p* < 0.05 and \*\*\**p* < 0.001.

Intuitively, the role of animals in transmitting CCHFV will largely depend on the level of viremia during infection, and only viremia above a certain threshold level will be sufficiently infectious. It is noteworthy, however, that the magnitude of viremia, or viral (RNA) load, did not always correlate with that of infectious virus. Three animals yielded successful virus isolation at the time points with peak viral load, whereas in Sheep 21-02 we were only able to isolate infectious virus from earlier time points with lower viral loads (2 DPI and 4 DPI), but not from the time point with peak viral load (6 DPI) (Figure 1). Similar discrepancy between the profiles of viral load and infectious virus titer was previously observed in CCHFV cell cultures [64]. The underlying mechanism is unknown, however this might be explained by the possibility that in some cases such as in Sheep 21-02, viral RNA could accumulate over time in the form of defective virus resulting from inactivation by the host immune response yet remaining to be cleared, or in the form of replication products including immature, partially assembled virions released from lysed host cells. Nevertheless, it should be noted that CCHFV infectivity appeared to be highly efficient, even at viral RNA concentrations as low as a few copies (cp)/µl. These were 0.6 and 3.5 cp/µl, respectively, on 2 DPI and 4 DPI in Sheep 21-02 (Figure 1), which are near the detection limit of modern real-time PCR methods. Therefore, a low-level or potentially undetectable viremia does not necessarily indicate the lack of infectious virus and does not rule out the capability of the animal host to spread the virus.

### Clinical signs, hematological features and biochemical characteristics

While no major disease was apparent (Table S1), all animals consistently demonstrated a significant increase in body temperatures following infection (Table S1 and Figure 2). We measured rectal temperatures and defined those above 39.9 °C as higher than normal body temperatures (fever) according to the literature [80, 81].

**Figure 2.**
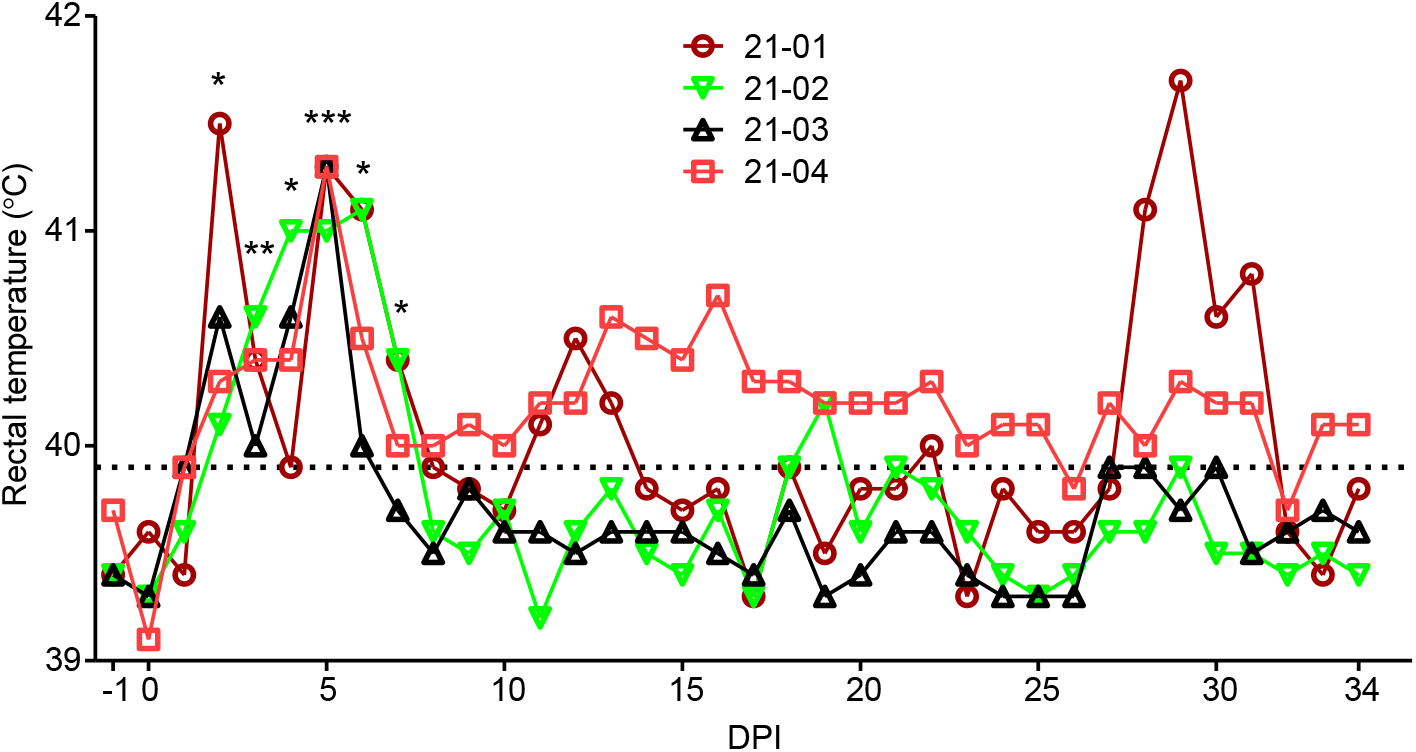
Fever in CCHFV-infected sheep. Line graph shows rectal temperature time course of each animal. Dash line indicates threshold temperature that defines a fever. Temperature significantly spiked following CCHFV infection as compared to the baseline (−1 DPI): \**p* < 0.05, \*\**p* < 0.01 and \*\*\**p* < 0.001.

Fever was the most dramatic during the viremic period, with peak temperatures (above 41 °C in all animals) largely occurring on or near 5 DPI (Table S1 and Figure 2), coinciding with peak viremia (Figure 1). Interestingly, elevated temperatures were also observed after the viremic period (Figure 2). It is unclear whether these resulted from a persistent effect of CCHFV infection; however, a prolonged fever in Sheep 21-04 and a late fever spike in Sheep 21-01 appeared to be associated with high levels of persistent viral RNA in tissues (see below).

Hematological changes in total white blood cells, monocytes, red blood cells or platelets were not found (Table S1). Consistent with the normal platelet parameters, prothrombin time and activated partial thromboplastin time also showed no change (Table S1). However, all animals demonstrated a sharp fall in neutrophil count and percentage (relative to total white blood cells) in the time window of 1 DPI to 3 DPI, followed by a rebound to levels higher than the baseline and return to near the baseline at later time points (Figure 3A and B). In addition, a significant spike in lymphocyte count was detected following peak viremia, at time points ranging from 7 DPI to 9 DPI (Figure 3C), possibly reflecting the expansion of lymphocytes during advancing adaptive immune responses. The maintenance and expansion of lymphocytes in sheep represents a distinctive feature as lymphopenia is a hallmark of severe CCHF in humans, cynomolgus macaques and immunodeficient mice [32, 63, 82].

**Figure 3.**
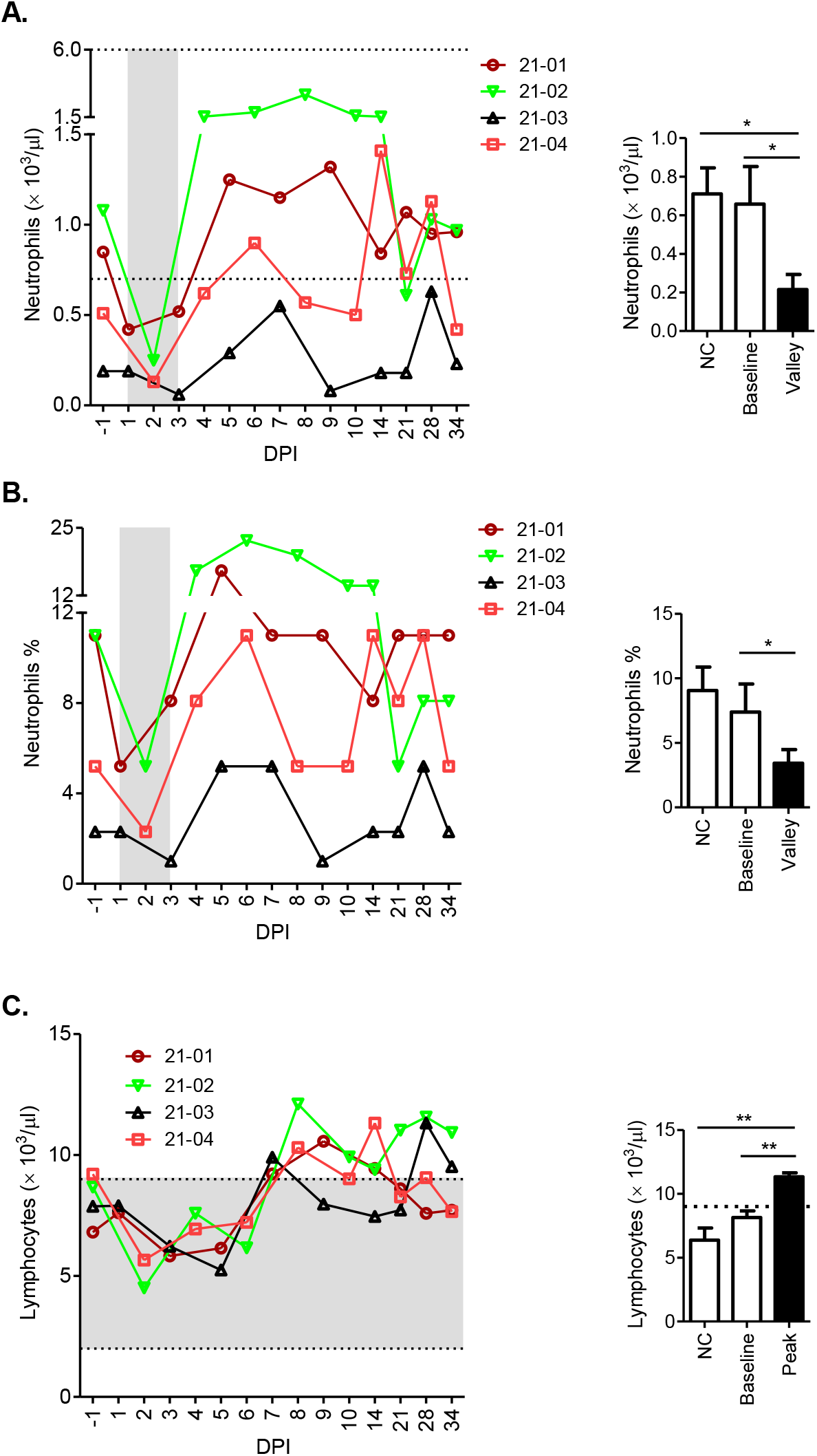
Hematological changes following CCHFV infection. Data on neutrophil count (A), neutrophil % within the leucocyte population (B) and lymphocyte count (C) are presented in graph formats similar to those in Figure 1. Grey columns in A and B highlight window of an early fall of neutrophil count or percentage to a bottom level consistently found in all animals. For contrasting purpose, double dash lines in A and C indicate the upper and lower limits of normal parameter ranges commonly observed for sheep in the literature, and single dash line in C indicates the upper limit.

The role of neutrophils in the control or pathogenesis of CCHFV infection is poorly understood [53, 83]. Recent research has revealed a large functional versatility of neutrophils as mediators of immune response and inflammation including antiviral responses that limit viral replication and expansion, beyond their traditionally recognized role in simply killing invading bacteria. Neutrophils have also been implicated in pathological processes such as dysregulated inflammation, acute organ injury, and coagulation abnormalities [84]. Activated neutrophils produce pro-inflammatory cytokines such as IL-8, the prototype and most potent neutrophil-attracting and neutrophil-activating cytokine [85], and TNF-α, which can in turn activate more neutrophils and other immune cells [84, 86]. Studies suggested that a positive feedback loop of systemic and neutrophil autocrine IL-8 production and a feed-forward cascade involving neutrophils and pro-inflammatory cytokines (including TNF-α) contribute to the severity of COVID-19 and influenza diseases, respectively [87, 88]. With potential relevance to these findings, lethal CCHFV infection in interferon α/β receptor knockout mice was featured by neutrophil infiltrations found in the liver and spleen [83]. We hypothesize that neutrophils may exert a pathological effect on susceptible hosts toward fatal CCHF disease. Sheep may evade or counter such effect by restricting the recruitment of neutrophils into the blood stream. This appeared to occur as an early response to CCHFV infection, as observed on 1 DPI – 3 DPI (Figure 3A and B). The restriction of recruitment may be through decreased mobilization of neutrophils from bone marrow into blood, increased storage or retention in the bone marrow or/and diminished production in the bone marrow [89]. Notably, the serum level of neutrophil chemoattractant IL-8 demonstrated a dramatic drop (see below) corresponding to the fall of neutrophils in the blood.

A series of blood biochemical parameters (https://www.zoetis.es/_locale-assets/spc/rotor-vs2-comprehensive-diagnostic-profile.pdf) were utilized mainly as markers for potential disease in liver and kidney, which are typical targets by fatal CCHFV infection. Of the two most widely used liver disease markers for severe and fatal human CCHF, our analytical kit included ALT, without AST available. Significant changes were not found in albumin, alkaline phosphatase, ALT, bilirubin, calcium, phosphorus, creatinine, sodium, potassium, total protein or globulin (Table S1). These results suggest that there was no major injury to the liver or kidney. However, blood urea nitrogen (BUN) and calcium levels in all the animals demonstrated a consistent and significant drop in early time windows of 1-2 DPI and 1-3 DPI, respectively, although with rebound and fluctuation at later time points (Figure 4A and B). In addition, all animals showed a trend of increase in blood glucose levels following infection, with subsequent decline or fluctuation (Figure 4C).

**Figure 4.**
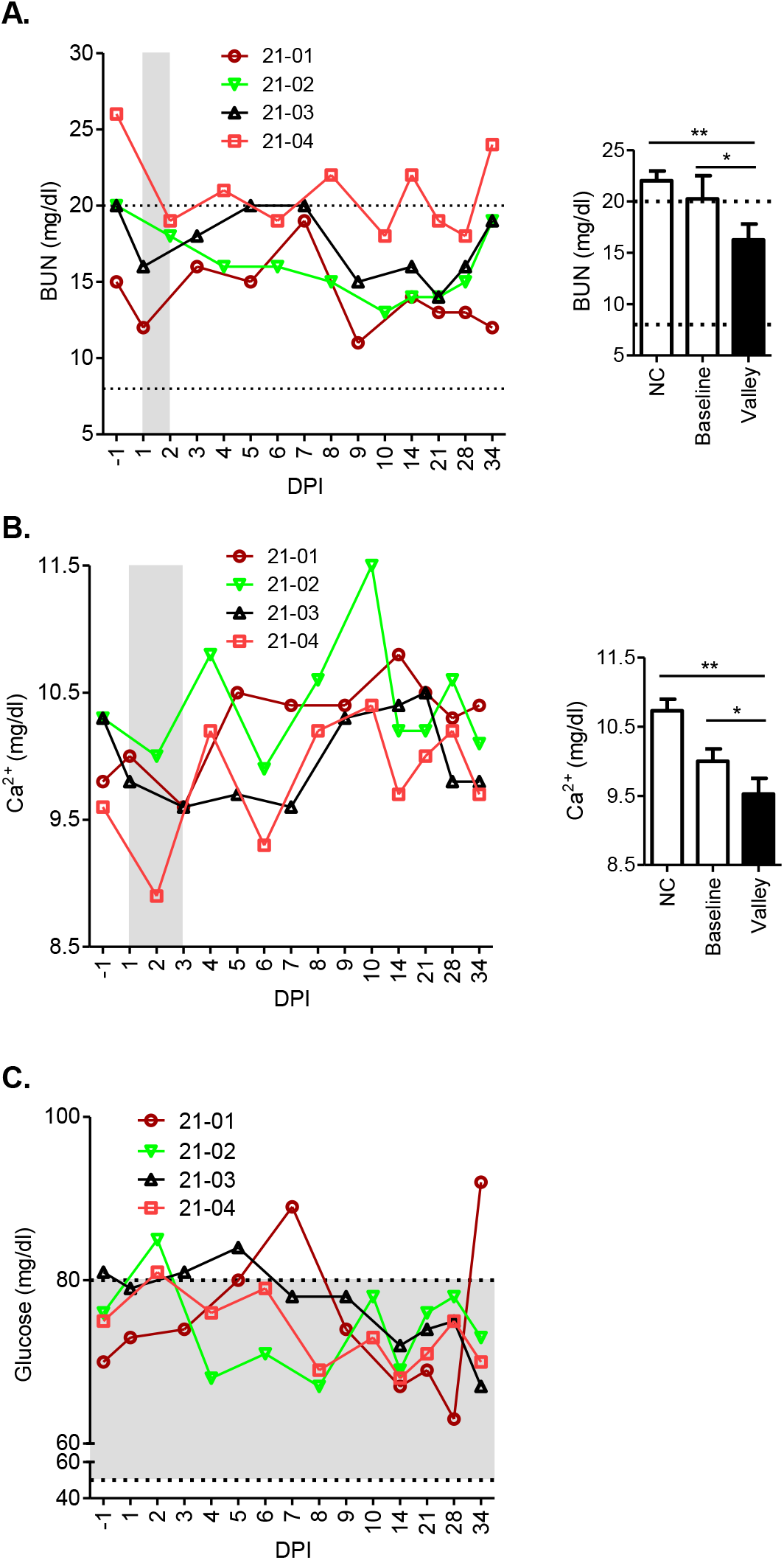
Blood chemistry changes following CCHFV infection. Data are presented in graph formats similar to those in Figure 3. BUN, blood urea nitrogen.

A major known cause of low BUN levels is liver disease [90–92]. Low blood calcium levels is a marker for kidney disease or worsening kidney function even when it is within the normal range or only mildly decreased [93, 94]. High blood glucose levels can result from increased hepatic glucose production and decreased renal glucose uptake caused by attenuated response to insulin in these organs under disease condition [95, 96]. Therefore, the changes in BUN, calcium and glucose levels together appeared to suggest a transient and minor impairment in certain biochemical functions of the liver and kidney. In CCHFV-infected African sheep, a slight but significant increase in AST was found, but no change in ALT was observed [79]. These results together with ours suggest that CCHFV infection in sheep may differentially affect some but not all disease markers since the effect of infection is not as major and extensive as in humans. However, the exact impact of CCHFV infection, especially potential long-term sequelae, on animal health status and production remains to be determined, as this area has never been covered by in-depth investigations, and variations in the biology caused by CCHFV infection among the animal population may have gone unrecognized.

### Immunological findings

All animals developed antibodies in the serum to CCHFV glycoprotein (Gn) and nucleoprotein (Figure 5A and B). Anti-Gn IgG antibodies were first detected on 7 or 8 DPI and quickly spiked on 8 – 10 DPI. Interestingly, a drop of antibody levels (as detected on 14 and 21 DPI) was followed by a rebound (28 and 34 DPI), which was consistently seen in all the animals (Figure 5A). This is reminiscent of the boost of a receding antibody response by a recurring antigen exposure. Anti-nucleoprotein IgG antibodies were initially detected on 4 or 5 DPI and demonstrated a gradual and continuous increase, except at late time points a trend of decrease followed by a rebound, similar to the observed pattern in anti-Gn antibodies (Figure 5B). The trend of decrease was perceivable at time points ranging from 10 to 21 DPI but to a lesser extent than in anti-Gn antibodies. The subsequent rebound, observed on 28 and 34 DPI, was characterized by a further elevation above the levels prior to the decrease (Figure 5B). These findings imply a recurrence of CCHFV antigen long after the resolution of viremia, which appears to be consistent with a tissue persistence of CCHFV RNA as described later. Virus neutralization activity was detectable at nearly all time points on or after 6 DPI (Figure 5C). Together, the antibody data indicate that CCHFV-infected sheep mount a quick antibody response, in contrast to the lack of antibody response in fatal human cases or immunodeficient mice, which is a strong predictor of poor outcomes.

**Figure 5.**
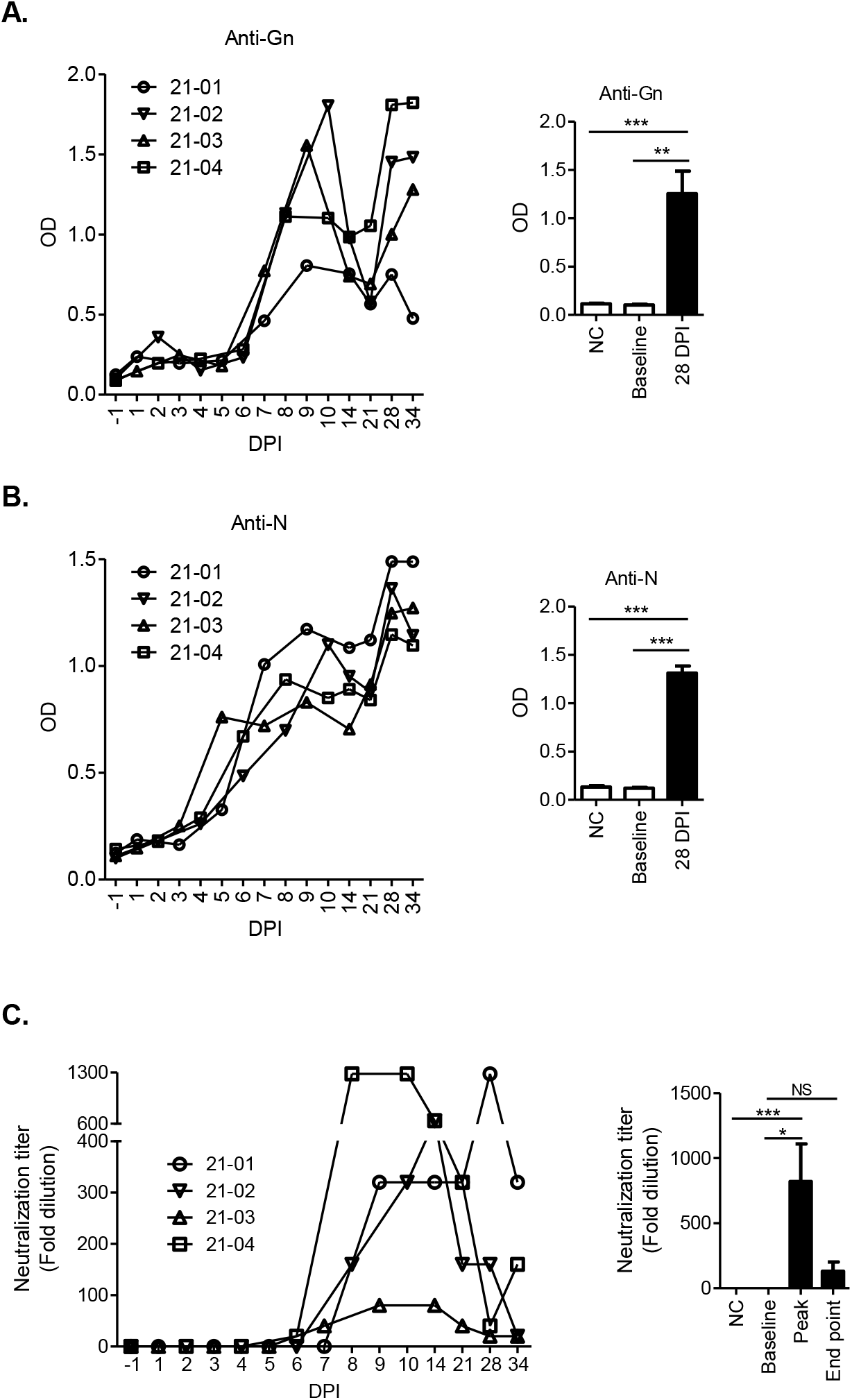
Antibody responses in CCHFV-infected sheep. **A and B.** Serum levels of anti-CCHFV Gn or nucleoprotein (N) IgG antibodies were measured by ELISA. OD, optical density. **C.** Neutralizing antibody titers in the serum were measured by virus neutralization test.

Serum levels of 14 cytokines were quantified (Figures 6-8). According to traditional classifications, IFN-γ, IP-10, IL-1α, MIP-1α, IL-8, VEGF-A, IL-1β, IL-17A, MIP-1β and TNF-α are pro-inflammatory cytokines; IL-6 is typically regarded as a pro-inflammatory cytokine but has also demonstrated anti-inflammatory properties; and IL-10, IL-4 and IL-36Ra are anti-inflammatory cytokines [53, 97–103].

**Figure 6.**
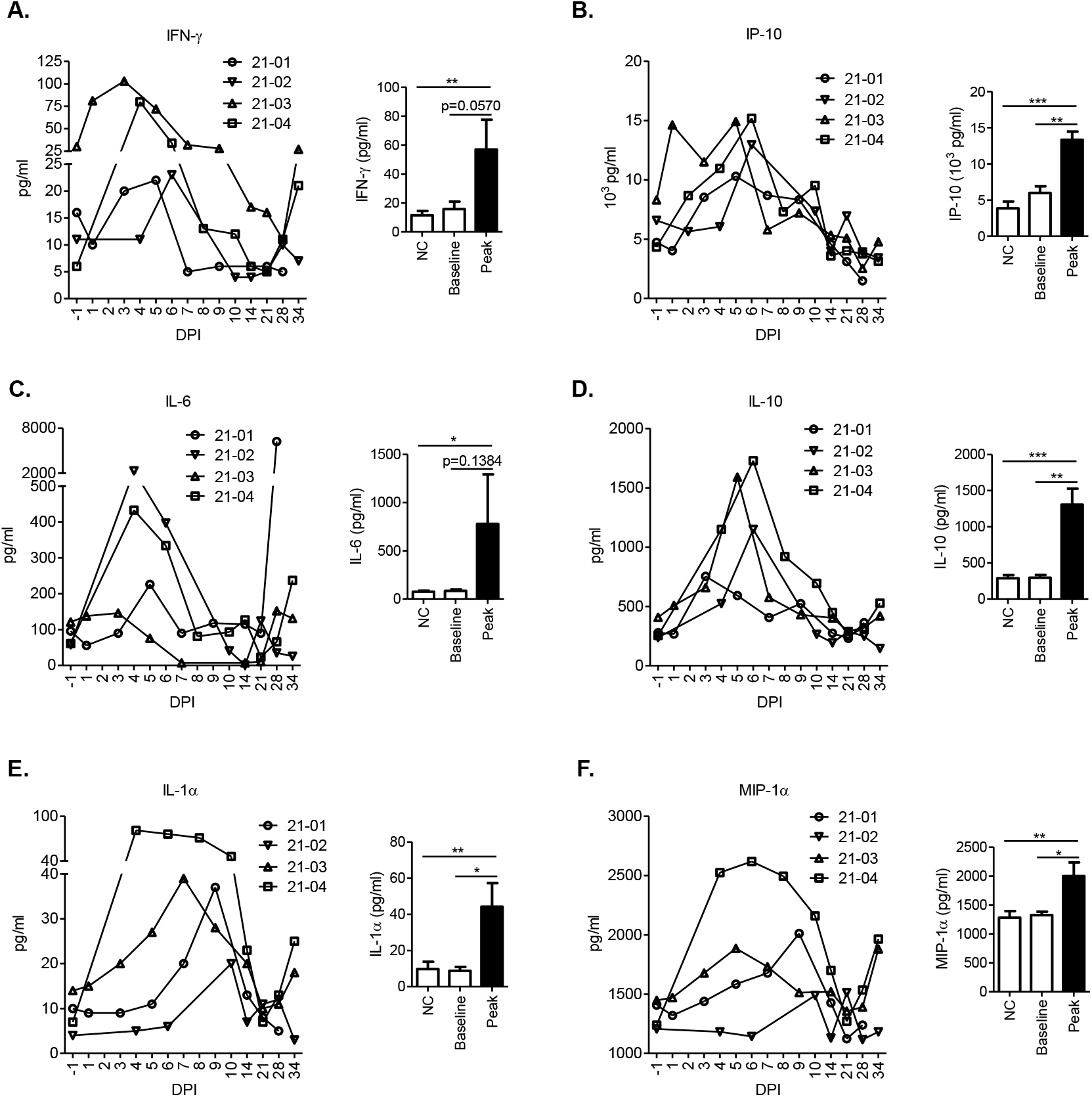
Cytokines that demonstrated increases following CCHFV infection.

**Figure 7.**
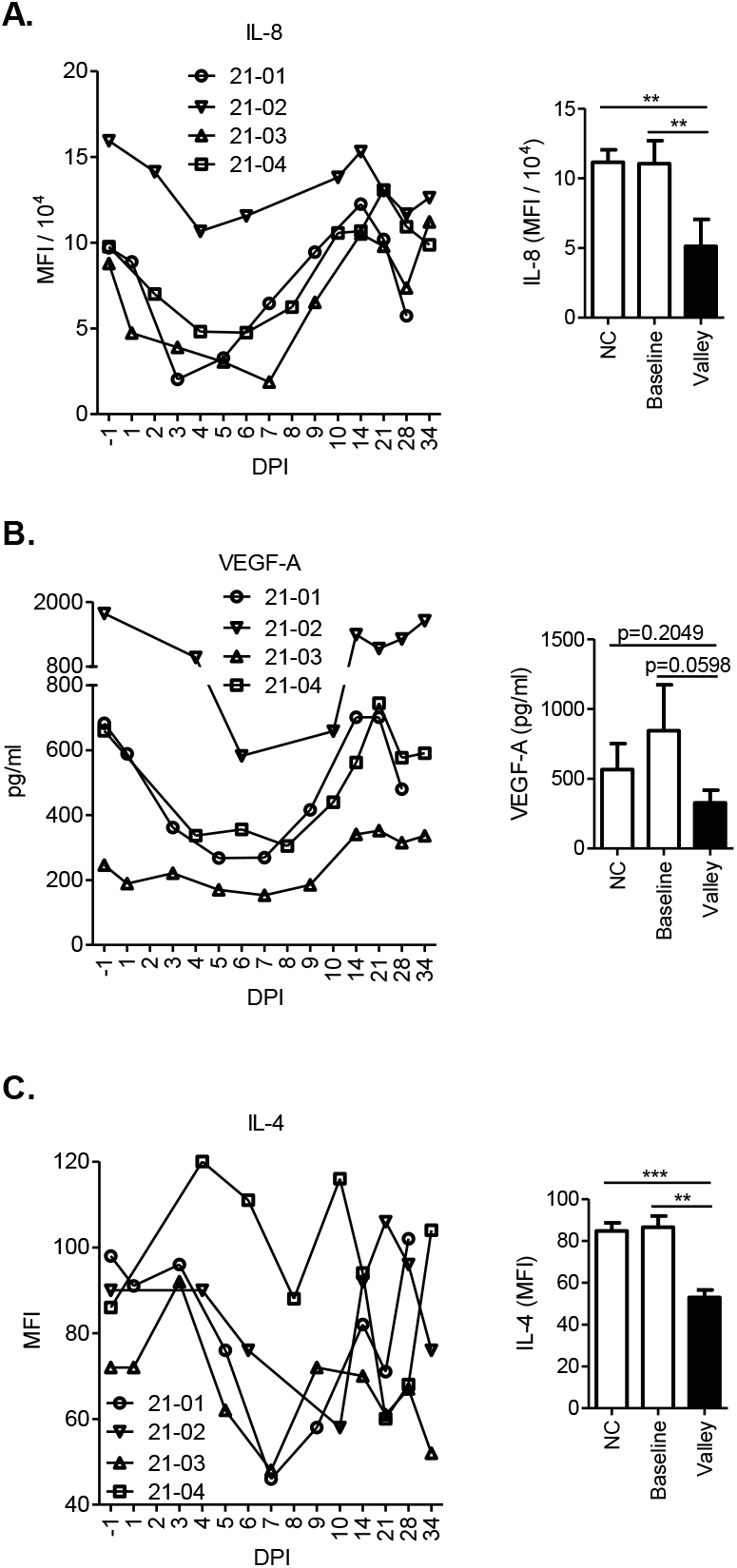
Cytokines that demonstrated decreases following CCHFV infection.

**Figure 8.**
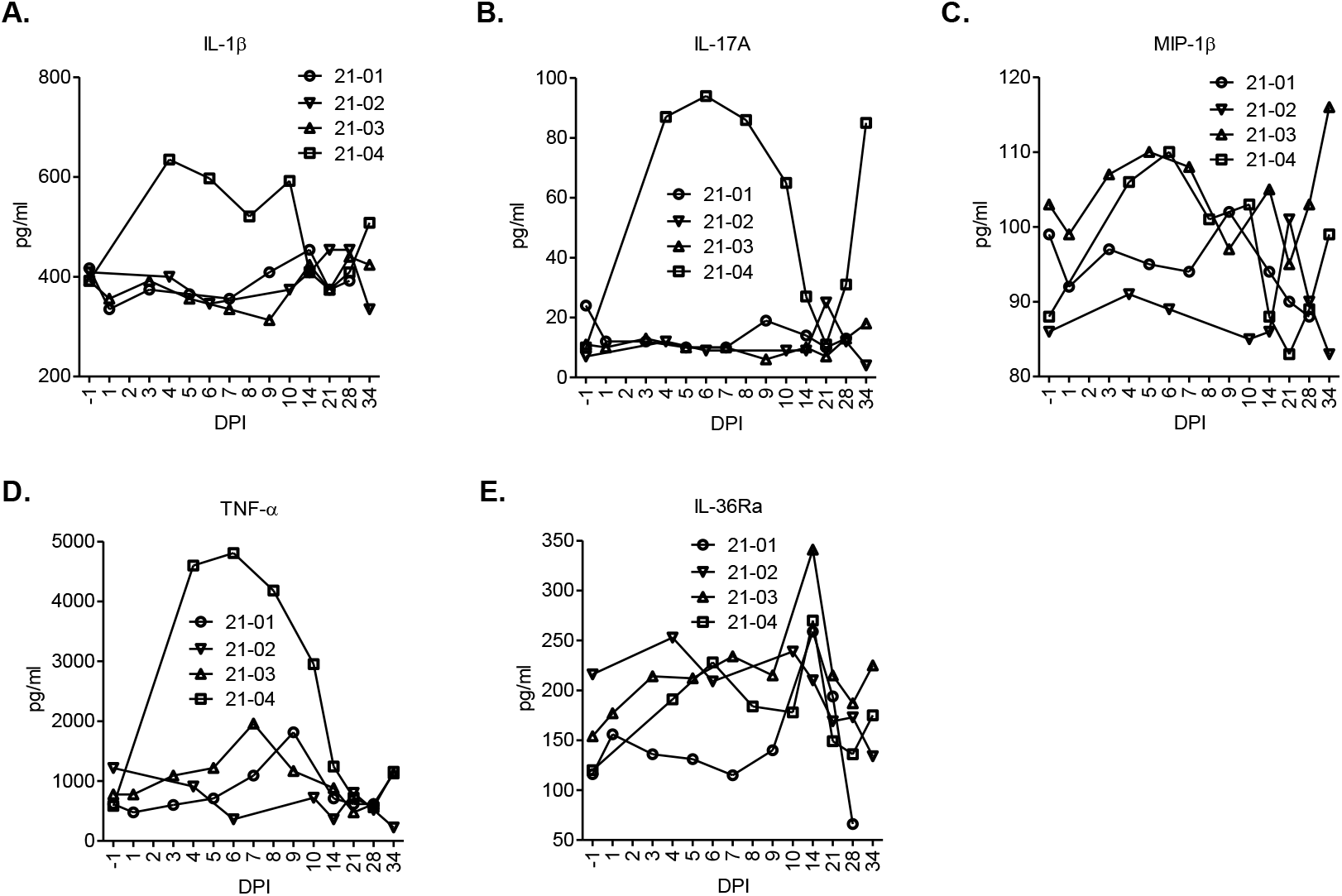
Cytokines that did not show consistent and significant changes following CCHFV infection.

In response to CCHFV infection, IFN-γ, IP-10, IL-6 and IL-10 exhibited significant increases largely coinciding with the viremic period followed by declines after peak viremia (Figure 6A-D). Increases were also found in IL-1α and MIP-1α, but with delayed kinetics extended into post-viremic time points (Figure 6E and F). A dramatic decrease in IL-8 was observed since 1 or 2 DPI, with the lowest IL-8 levels falling on time points around peak viremia. This was followed by a rebound that reached baseline levels by approximately 21 DPI (Figure 7A). A trend of similar decrease (in all animals) was found in VEGF-A, while it did not reach statistical significance (Figure 7B). Following an early trend of slight increase, IL-4 demonstrated a sharp drop after 3 or 4 DPI, with the lowest levels reached at time points shortly after peak viremia (7, 8 or 10 DPI) and followed by rebound later (Figure 7C). No changes consistent among all the animals were found in IL-1β, IL-17A, MIP-1β, TNF-α or IL-36Ra (Figure 8A-E). A trend of IL-36Ra spike (in three of the four animals), however, was noticed at 14 DPI (Figure 8E), overlapping with the timing when the decreases in anti-Gn and anti-nucleoprotein antibodies as described above started to be observed (Figure 5A and B).

It has been proposed that CCHF pathogenesis in humans could be a result from massive, dysregulated inflammatory responses, possibly in combination with direct tissue injury by the virus, and intensely increased pro-inflammatory cytokines (cytokine storm) may mediate vascular dysfunction, DIC, organ failure and shock [25, 28, 104, 105]. Out of a number of cytokines that have been assessed so far, increases in TNF-α, IL-8, IL-6, IL-1β, IP-10, VEGF-A and IFN-γ were associated with poor outcomes [28, 104, 106–110]. Of those, TNF-α has been the most frequently reported predictive indicator of mortality. Recent studies have made progress toward understanding whether these correlations reflect causal, rather than purely coincidental, relationships. Blocking TNF-α signaling with TNF-α receptor knockout or a TNF-α neutralizing antibody afforded survival advantage in mouse models [58]. Furthermore, TNF-α derived from CCHFV-infected monocyte-derived dendritic cells mediated endothelial cell activation [111]. It was also known that TNF-α, IL-8, VEGF-A, IL-1β and IL-4 increase endothelial permeability [112–124]. These are of significant mechanistic relevance as endothelial dysfunction and damage with leakage of erythrocytes and plasma through the vasculature into tissues is a hallmark of CCHF pathology. Endothelial damage contributes to coagulopathy by stimulating platelet aggregation and degranulation, with subsequent activation of the intrinsic coagulation cascade, leading to clotting factor deficiency and consequently hemorrhages [24, 25].

In contrast to fatally infected humans, CCHFV-infected sheep were able to down-regulate these cytokines (IL-8, VEGF-A and IL-4; Figure 7) or limit their increase (TNF-α and IL-1β; Figure 8), which may serve as a defense mechanism against vascular dysfunction. The reduction of IL-8 in sheep may also be part of an inhibitory circuit restricting pathogenic recruitment of neutrophils as mentioned above. It should be noted that these possibilities remain to be tested in further experiments and other possibilities exist. Differential responses were similarly observed between interferon-α/β receptor knock-out mice and sheep. In these mice, lethal CCHFV infection exhibited significantly elevated levels of TNF-α, IL-1β, IL-4 and IL-17A [83], whereas these fell or failed to increase in sheep. IL-17A was suggested in a previous study to synergize with TNF-α in neutrophil mobilization [125], which appeared to be limited in sheep.

The increases in other cytokines in CCHFV-infected sheep, along with the decreases or lack of significant changes in those discussed above, appeared to coincide with an induced anti-viral state with sharp reduction in viremia. Some increased cytokines may be part of an anti-viral response. Of these, IFN-γ was found to be critical for survival following CCHFV infection in interferon-α/β receptor knock-out mice [126]. Overall, the inflammatory cytokine responses were executed in sheep in a balanced manner as the effective control and resolution of viremia and the lack of remarkable inflammatory pathology were both achieved. The major changes in cytokine levels mostly corresponded to a limited window of acute viremia followed by their return to normal ranges. The timing, intensity and duration of these responses appeared to be within the ranges well tolerated by the host. As part of variations among individual animals, however, higher levels of some cytokines were noticed in sheep 21-04 including IL-1α, MIP-1α, IL-4, IL-1β, IL-17A and TNF-α (Figures 6-8). This was accompanied by a prolonged fever (Figure 2) and an extremely high level of viral RNA persistence in tissues (see below), while no major changes in the infection outcome were apparent. These results imply a complex nature of the biology underlying the lack of major disease, which was likely contributed to by the combination of multiple control mechanisms. Nevertheless, the immunological responses identified here especially those that differentiate between disease resistance and susceptibility host phenotypes provide promising targets for further mechanistic studies.

### Viral shedding, dissemination and tissue persistence

The virus spread beyond the blood in all animals. Nasal, oral and rectal shedding of viral RNA was detected in swab elutes from several time points of the viremic period, ranging from 3 DPI to 6 DPI (Table 1). We analyzed tissues collected from necropsies on 34 DPI, the study end point, anticipating that no viral presence would be detected since the animals appeared to be able to clear the virus early based on the resolution of viremia by 6 DPI (Figure 1). To our surprise, however, viral RNA was detected extensively in multiple types of lymph nodes (in all animals) as well as in the liver and spleen (in all animals except Sheep 21-04), and occasionally in other organ/tissue types including the ileum, adrenal gland, lung and potentially cerebrospinal fluid (Table 2). Among these, Sheep 21-04 and 21-01 demonstrated exceptionally high viral RNA levels in inguinal lymph nodes (11082.04 cp/µl) and deep cervical lymph nodes (2514.43 cp/µl), respectively (Table 2). These results indicate a widespread viral dissemination in the host and suggest a long-term persistence in tissues. It remains to be determined, however, whether functional virions are harbored or could be potentially produced in these tissues. Our virus isolation attempt, by applying tissue homogenates to SW-13 cell cultures, did not recover infectious virus. This could result from the limitation of the isolation conditions or the absence of readily packaged infectious virus.

**Table 1.** Viral RNA shedding.

| Sheep ID | Shedding source | CCHFV RNA (cp/μl) |  |  |  |  |  |  |  |  |  |  |  |  |  |  |
| --- | --- | --- | --- | --- | --- | --- | --- | --- | --- | --- | --- | --- | --- | --- | --- | --- |
|  |  | -1 DPI | 1 DPI | 2 DPI | 3 DPI | 4 DPI | 5 DPI | 6 DPI | 7 DPI | 8 DPI | 9 DPI | 10 DPI | 14 DPI | 21 DPI | 28 DPI | 34 DPI |
| 21-/01 | Nasal | -/- | -/- |  | -/- |  | -/- |  | -/- |  | -/- |  | -/- | -/- | -/- | -/- |
|  | Oral | -/- | -/- |  | -/- |  | -/- |  | -/- |  | -/- |  | -/- | -/- | -/- | -/- |
|  | Rectal | -/- | -/- |  | **2.37** |  | **127.27** |  | -/- |  | -/- |  | -/- | -/- | -/- | -/- |
| 21-/02 | Nasal | -/- |  | -/- |  | -/- |  | -/- |  | -/- |  | -/- | -/- | -/- | -/- | -/- |
|  | Oral | -/- |  | -/- |  | -/- |  | **1.14** |  | -/- |  | -/- | -/- | -/- | -/- | -/- |
|  | Rectal | -/- |  | -/- |  | -/- |  | **8.85** |  | -/- |  | -/- | -/- | -/- | -/- | -/- |
| 21-/03 | Nasal | -/- | -/- |  | **41.81** |  | **78.12** |  | -/- |  | -/- |  | -/- | -/- | -/- | -/- |
|  | Oral | -/- | -/- |  | -/- |  | **1.10** |  | -/- |  | -/- |  | -/- | -/- | -/- | -/- |
|  | Rectal | -/- | -/- |  | -/- |  | **4.39/-** |  | -/- |  | -/- |  | -/- | -/- | -/- | -/- |
| 21-/04 | Nasal | -/- |  | -/- |  | -/- |  | **3.16** |  | -/- |  | -/- | -/- | -/- | -/- | -/- |
|  | Oral | -/- |  | -/- |  | **2.28** |  | **23.14** |  | -/- |  | -/- | -/- | -/- | -/- | -/- |
|  | Rectal | -/- |  | -/- |  | -/- |  | -/- |  | -/- |  | -/- | -/- | -/- | -/- | -/- |

CCHFV RNA in swab elutes was quantified by RT-qPCR. Mean value of duplicate samples is shown except "-/-" meaning both duplicate samples were negative in PCR and "#/-" meaning one of the duplicates was positive while the other negative. Bold text indicates result with clinical significance. Empty spot indicates that sample was not collected due to sampling on alternate days.

**Table 2.**
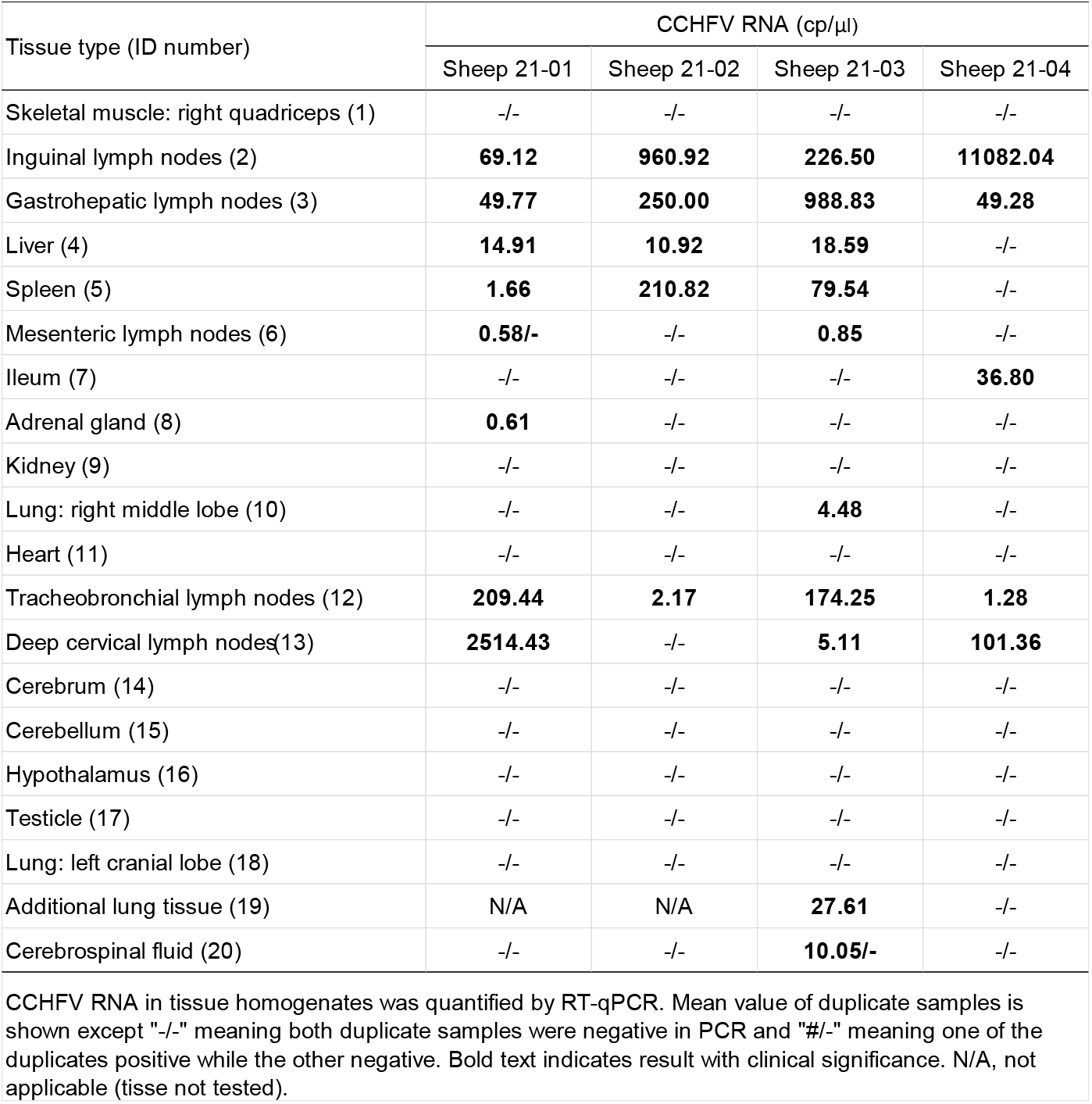
Viral RNA in tissues from 34 DPI.

| Tissue type (ID number) | CCHFV RNA (cp/μl) |  |  |  |
| --- | --- | --- | --- | --- |
|  | Sheep 21-01 | Sheep 21-02 | Sheep 21-03 | Sheep 21-04 |
| Skeletal muscle: right quadriceps (1) | -/- | -/- | -/- | -/- |
| Inguinal lymph nodes (2) | **69.12** | **960.92** | **226.50** | **11082.04** |
| Gastrohepatic lymph nodes (3) | **49.77** | **250.00** | **988.83** | **49.28** |
| Liver (4) | **14.91** | **10.92** | **18.59** | -/- |
| Spleen (5) | **1.66** | **210.82** | **79.54** | -/- |
| Mesenteric lymph nodes (6) | **0.58/-** | -/- | **0.85** | -/- |
| Ileum (7) | -/- | -/- | -/- | **36.80** |
| Adrenal gland (8) | **0.61** | -/- | -/- | -/- |
| Kidney (9) | -/- | -/- | -/- | -/- |
| Lung: right middle lobe (10) | -/- | -/- | **4.48** | -/- |
| Heart (11) | -/- | -/- | -/- | -/- |
| Tracheobronchial lymph nodes (12) | **209.44** | **2.17** | **174.25** | **1.28** |
| Deep cervical lymph nodes (13) | **2514.43** | -/- | **5.11** | **101.36** |
| Cerebrum (14) | -/- | -/- | -/- | -/- |
| Cerebellum (15) | -/- | -/- | -/- | -/- |
| Hypothalamus (16) | -/- | -/- | -/- | -/- |
| Testicle (17) | -/- | -/- | -/- | -/- |
| Lung: left cranial lobe (18) | -/- | -/- | -/- | -/- |
| Additional lung tissue (19) | N/A | N/A | **27.61** | -/- |
| Cerebrospinal fluid (20) | -/- | -/- | **10.05/-** | -/- |

CCHFV RNA in tissue homogenates was quantified by RT-qPCR. Mean value of duplicate samples is shown except "-/-" meaning both duplicate samples were negative in PCR and "#/-" meaning one of the duplicates positive while the other negative. Bold text indicates result with clinical significance. N/A, not applicable (tissue not tested).

Similar phenomena of viral RNA persistence after the clearance of acute infection have been emerging in other non-retroviral RNA viruses including Ebola virus, Marburg virus, Zika virus and severe acute respiratory syndrome coronavirus 2 (SARS-CoV-2). These have been linked to recurring, chronic or progressive post-viral disease or syndromes and late transmissions of infectious virus that can spark new outbreaks [127, 128]. It remains unknown in what forms and by what mechanisms the viral RNA persists. A lack of culturable virus together with the known susceptibility of RNA to degradation points to the assumption that the viral sequences detected by RT-PCR could be from fragmented RNA [127, 129]. However, dormant viral forms capable of reactivation, possibly with full-length RNA, have been suggested by the recrudescence of viral transcription and protein synthesis or infectious virus production [127, 130–137]. It has been proposed that latent viral RNA may be protected in the cytoplasm of infected cells as ribonucleoprotein complexes or by association with membrane structures. Viral and host factors may suppress the production of infectious virions facilitating the survival of both the host cell and viral RNA against immune recognition with subsequent clearance [127]. Awakening from dormancy can occur when immune control is relaxed or in response to certain stimuli [127, 130–134].

Whether a similar dormant form of CCHFV persisted in the sheep is an open question. The antibody waves suggestive of recurring production of CCHFV protein antigens (Figure 5) appear to be consistent with this possibility. It should be noted that RNA persistence previously observed in other viruses was in the context of pathogenic infection, whereas CCHFV does not cause prominent disease in sheep and a widespread viral RNA persistence is less anticipated. Thus, this unique finding extends the known spectrum of types of infections with viral RNA persistence. Long-term health effects of CCHFV RNA persistence in sheep (notably related to the liver and the lymphatic system), however, should only be excluded by extended experimental studies, as these effects could potentially go unrecognized as seemingly non-specific variations in health status among the animal population. It must be noted that the highest levels of viral RNA persistence (Table 2) appeared to correlate with a prolonged fever and higher levels of inflammatory cytokines in Sheep 21-04 (Figures 2 and 6-8) and a late fever spike in Sheep 21-01 (Figure 2).

CCHFV RNA in the lymphoid organs may serve as sources of persistent antigenic stimulations directly within the immune system, as hinted by the anti-Gc and nucleoprotein antibody waves (Figure 5). Persistent immune activation has been implicated in immune exhaustion and increased re-infection by SARS-CoV-2 [129]. On the other hand, continued availability of antigenic boost is believed to be beneficial to the host for replenishing immunity such as in measles virus [138, 139]. These different possibilities warrant longer-term follow-up investigations in CCHFV-infected sheep.

It is noteworthy that windows of immune relaxation could allow the occasional release of infectious virus from latent reservoir reactivation as observed in Ebola, Zika and measles viruses, with relevance to late transmissions [133–137, 140–142]. Infectious virus potentially inducible from dormant CCHFV might be isolated with improved methods, which could be based on co-cultivation of tissues harboring persisting CCHFV RNA (as opposed to tissue homogenates) with susceptible cells, possibly in combination with immunosuppressive stimuli such as cyclophosphamide [133, 134, 140–142].

Epidemiological surveillance of CCHFV infections that have occurred in animals commonly depends on serological tests for CCHFV antibodies in conjunction with the ease of blood sampling. The use of the more sensitive and specific molecular methods, notably based on real-time PCR, has been limited by the consideration that viral RNA is only present in the blood for a short window during active replication. However, the findings of CCHFV RNA persistence suggest that PCR analysis of tissue RNA can be employed as an additional method to assess CCHFV infection in sheep (or possibly other animals to be found with CCHFV RNA persistence). Tissues collected during livestock slaughtering, for example, can serve as materials for testing. The PCR surveillance may complement serosurveys by catching cases corresponding to a waning phase of antibody responses, while it will provide information on the prevalence of CCHFV RNA persistence in animal populations.

## Conclusions

CCHFV Kosovo Hoti-infected sheep developed a viremia in the absence of prominent clinical signs, confirming observations from past studies [29]. A cryptic CCHFV infection in livestock could pose great risk to public health due to unexpected transmission, especially in new regions where viral prevalence has not yet been noticed. In addition, although no major disease was manifested in infected animals, illness may be present but unrecognized. Markers for potential impairment in liver and kidney functions and viral RNA persistence in the liver, spleen, lymph nodes and some other types of organs/tissues, with a prolonged fever or late fever spike associated with high levels of viral RNA persistence, do advise possible impact of hidden CCHFV disease on animal health status and production levels, which has never been covered in experimental studies.

The differential outcomes of CCHFV infection in sheep and humans provide an opportunity for investigations into the host factors that control disease. Distinctive immune responses were identified in sheep that distinguish their subclinical infection from fatal infection in humans. CCHFV-infected sheep were able to maintain and expand lymphocytes and develop quick antibody and cytokine responses associated with a rapid resolution of viremia. Notably, an early restriction of neutrophil recruitment and IL-8 levels may prevent pathogenic neutrophil infiltrations, and multiple cytokines with known roles in endothelial damage were found to be down-regulated or limited from increase, which may avoid vascular dysfunction and subsequent progression to severe disease. Genetic or pharmaceutical targeting of these distinguishing responses will determine whether they act as protective mechanisms against disease in sheep. Resulting knowledge will inform medical countermeasures that promote beneficial immune responses while limiting immunopathology.

The current knowledge regarding the host determinants of CCHFV infection outcomes has been hindered by the lack of ideal animal models. Studies have largely been limited to and focused on immunodeficient mice, as the field has been struggling to find an immunocompetent CCHF disease model. The mouse models have greatly contributed to our understanding of CCHFV pathogenesis but have limitations. These mice have incomplete and altered immune systems and type I interferon deficiency impacts both innate and adaptive immunity, which could lead to confounded findings or missing insights into natural responses that could otherwise be obtained in immunocompetent hosts. Indeed, the importance of lessons learned from natural immunity has been demonstrated by the advances from other viral disease models [143–145]. Although comparative studies in the context of immunocompetent hosts may be considered in human patients with different outcomes of CCHFV infection, in general they could not be conducted in-depth experimentally with major genetic or therapeutic manipulations. Instead, CCHFV infection in sheep can serve as a valuable immunocompetent animal model for studying host factors involved in pathogenesis and disease outcome, supplementing the immunodeficient mouse infection models.

This study brings pioneering findings of extensive CCHFV dissemination, viral shedding and viral RNA persistence in tissues in the context of immunocompetent animal hosts. Viral spread beyond the viremia presents additional sources of potential viral transmission, while this study has also confirmed the previously recognized role of blood based on past experimental data. Together, these findings support public health education and measures aimed to prevent or reduce the risk of acquiring CCHFV infection from infected animals, apart from infected ticks. Professionals at high risk such as farmers, slaughterhouse workers, veterinarians and stockmen should be made aware of these potential sources of infection from animals. Under the One Health approach, tick control has been implemented by the use of acaricides, which can be practically difficult under extensive farming conditions [48]. Alternatively, veterinary vaccines that block CCHFV replication hold promise to break animal host-mediated viral transmission chains to ticks and humans. In this regard, CCHFV Kosovo Hoti infection in sheep represents a valuable livestock model for the development of such vaccines, where vaccine effects on viremia and viral dissemination, shedding and tissue persistence can be tested. In addition, with CCHFV RNA long-term presence in sheep tissues this infection model carries an outstanding potential for studying viral RNA persistence in the context of disease-resistant hosts. While the potential health impact of CCHFV RNA persistence remains to be clarified, it may likely promote a durable anti-viral immunity.

Concerning the major limitation of this study, it was a pilot test of viral dissemination and persistence in tissues for CCHFV in animals and thus included only one late time point of animal sacrifice for tissue collection following the 3Rs animal use ethics. An enhanced and extended time course investigation with serial animal sacrifices will next determine the temporal and spatial sequence of viral dissemination through the organs/tissues in the host. The availability of frequent and extended tissue time points will also facilitate extensive characterization of viral RNA persistence, including the determination of time points and locations at which readily infectious virus or inducible virus or antigens could be isolated or detected. This will require a larger number of animals, which is now justifiable based on the new findings from the current study.

In conclusion, our study reveals previously unrecognized aspects of CCHFV biology in animals, presents a significant value of the CCHFV sheep infection model for studying host factors controlling outcomes of infection, for testing veterinary vaccines and for characterizing viral RNA persistence, and encourages and provides perspectives for extended future studies based on the revisit of experimental infection in animals.

## Acknowledgements

We thank Cory Nakamura for performing sampling and necropsy, Yvon Deschambault for running hematology and blood chemistry tests of farm control sheep, Charles Lewis and Valerie Smid for providing technical advice and assistance, and Animal Care staff at the National Centre for Foreign Animal Diseases for animal care. This work was supported by a Canadian Safety and Security Program grant (CSSP-2018-CP-2341) and funding from Canadian Food Inspection Agency.

## Additional files

**Table S1. Clinical findings and blood results.** An excel spread sheet including four tabs each corresponding to one infected sheep.

